# Altered perceptual integration and learning in autism revealed by games inspired by rodent operant tasks

**DOI:** 10.1101/2025.05.06.652509

**Authors:** Sucheta Chakravarty, Quan Do, Yutong Li, Helen Tager-Flusberg, Joseph T. McGuire, Benjamin B. Scott

## Abstract

Altered perception is a key feature of autism spectrum disorder (ASD), yet its precise nature and variability across individuals remain unclear. We developed an online video game, inspired by rodent operant tasks, to assess visual evidence integration. The game employs nonverbal, reward-based training and a pulsed stimulus design for precise control of sensory input, enabling detailed individual-level behavioral analysis. It was playable by typically developing adolescents and their autistic siblings, who spanned the spectrum, including those with profound autism. Game performance correlated with standard ASD survey scores, with ASD participants exhibiting slower learning and altered perceptual integration. Behavioral data were well fit by computational models in which perceptual and learning deficits in ASD arose from increased noise in higher- order visual processing. Our findings reveal that deficits in perceptual integration are widespread across ASD, correlate with symptom severity in social and adaptive domains, and may arise from instability of sensory representations.

## Main

Atypical perceptual processing is considered a core feature of autism spectrum disorder (ASD; *Diagnostic and Statistical Manual of Mental Disorders*, 2013, Marco et al., 2011; Robertson & Baron-Cohen, 2017). However, the exact nature of perceptual alteration remains unclear (Balasco et al., 2020; Morimoto et al., 2021; Hadad & Yashar, 2022). Previous research suggests that sensory perception in ASD may be affected at multiple stages of processing (Simmons et al., 2009, Orefice, 2020; Orefice, 2019; Orefice et al., 2016). Psychophysical studies indicate that individuals with ASD perform similarly to typically developing (TD) controls on tests of simple detection or discrimination (Pellicano et al., 2005; Koh et al., 2010), whereas complex tasks, such as those requiring sensory evidence accumulation, reveal more significant impairments (Robertson et al., 2012). These findings have led to the idea that atypical sensory perception in ASD arises, in part, from deficits in higher-order cognitive processes, such as *perceptual integration* (Robertson & Baron-Cohen, 2017).

Perceptual integration is a key step in neural information processing when sensory stimuli are noisy or uncertain (Gold & Shadlen, 2007), or when the stimulus varies over time, as with human speech (Poeppel, 2003). Compared with TD individuals, individuals with ASD exhibit slower response times on perceptual decision-making tasks and lower accuracy rates on tasks that involve ambiguity or noise, suggesting that deficits in perceptual integration could contribute to altered sensory perception in ASD (Robertson & Baron-Cohen, 2017). Several models of altered sensory processing have been proposed to account for deficits in perceptual integration in ASD. Slower response times suggest elevated decision boundaries (Pirrone et al., 2020) or slower evidence accumulation rates (Robertson et al., 2012), whereas neurophysiological measurements and behavioral studies indicating variability in stimulus-evoked responses suggest increased noise in the perceptual process (Dinstein et al. 2012; Haigh, 2018). Whether the ASD phenotype involves differences in bottom-up sensory processing or top-down cognitive processing, or has multiple independent mechanisms that vary across individuals, is still unknown (Rosenberg et al., 2015; Balasco et al., 2020; Morimoto et al., 2021).

Large-scale psychophysical studies provide the quantitative rigor necessary to characterize sensory perception; however, several methodological challenges slow progress in this area in ASD (Hadad & Yashar, 2022). Previous studies have largely focused on small cohorts of ASD participants. Furthermore, these studies typically involve ASD populations matched with non-autistic comparison participants, and individuals with more limited language and intellectual disability have been underrepresented. The reliance on small sample sizes and reduced range of participants may contribute to conflicting results in perceptual studies of ASD and may limit our understanding of how perception varies across the spectrum (Koldewyn et al., 2010; Milne et al., 2002; Pellicano et al., 2005; Robertson et al., 2012; Price et al., 2012, Van der Hallen et al., 2019).

To address these challenges, we used a novel online video game to test perception and cognitive function across a large, heterogeneous sample of participants with ASD, including some with profound autism. The game is based on pulse-based psychophysical tasks originally developed in animal models to distinguish between alternative cognitive models of perceptual integration (Brunton et al., 2013; Scott et al., 2015; Kane et al., 2024), which have since been back-translated to humans (Do et al., 2023; Chakravarty et al., 2024). In the game, players make choices based on unreliable sensory information, integrating evidence over time to improve their scores. The game incorporates feedback-based training, inspired by animal studies, facilitating learning in individuals unable to follow detailed verbal instructions.

We measured gameplay online in a large cohort of adolescents with autism spectrum disorder (ASD) and their unaffected siblings. The games were playable by individuals across the spectrum, including individuals with profound autism. ASD participants displayed deficits in learning and perceptual processing. At the individual level, gameplay metrics correlated with symptom severity measured with traditional survey scores. Behavior modeling suggested that deficits in ASD participants were consistent with greater integration noise, which increased exponentially with the number of pulses. Together, these data support a model in which sensory and learning deficits in ASD arise, in part, from alterations in perceptual integration. Moreover, these results demonstrate the value of online video games with non-verbal training pipelines for efficient, high-throughput assessment of perceptual and cognitive functioning across ASD.

## Results

### Online video game for testing perceptual integration in ASD

We developed an online video game inspired by pulse-based evidence accumulation tasks previously used in rodents (**Fig. 1**). In our version, participants were presented with brief light pulses (flashes) to their left and right visual hemifields and were rewarded for choosing the side (left or right) with more flashes (**Fig. 1B**). The discrete nature of the pulses provides exquisite experimental control over the accumulation process. The number, duration, and timing of pulses can be varied per-trial to yield a rich dataset which facilitates model-based analysis of behavior to distinguish deficits in different stages of perceptual processing, from early sensory input to later stages (**Fig. 1A**).

**Figure 1.**
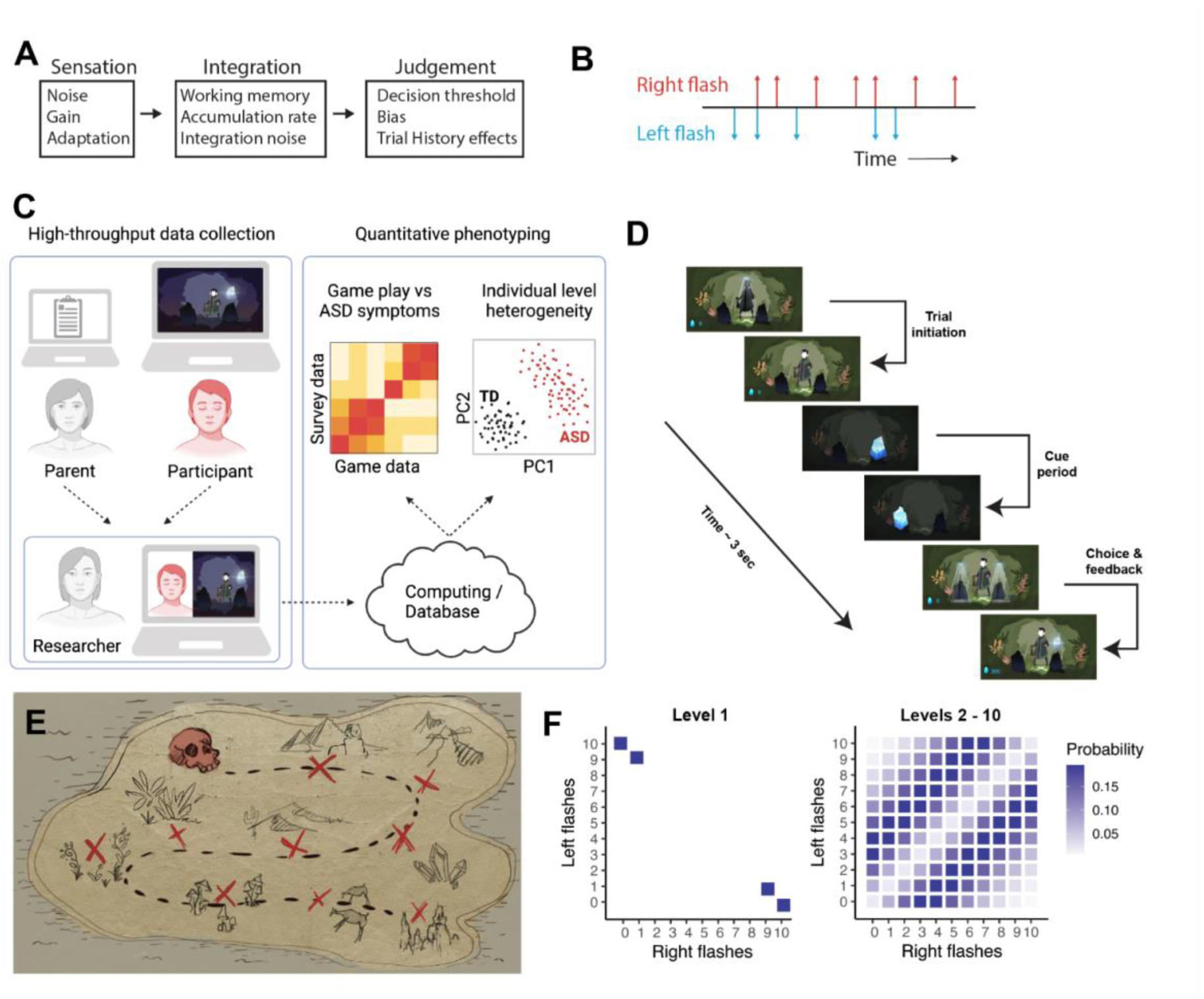
Overview of the approach. **A.** Atypical perceptual processing in ASD may be due to deficits in early sensory input or later processing stages, the exact nature of which remain to be understood. **B.** The pulse-based evidence accumulation task used brief light flashes presented to the left and right hemifields, at random time points. Participants were rewarded for choosing the side (left/right) with more flashes. **C.** We turned the pulse-based task into an online video game to collect high throughput data, which was used to compute multiple behavioral measures. This enabled more data-driven analysis of heterogeneous ASD phenotypes. **D.** Each trial in the game consisted of the following: initiation, cue period, and choice & feedback. Trials were initiated by clicking on the center; flashes (in the shape of gems) appeared during the cue period. After the cue period, the participant had to make a side choice and click on it; only correct choices ( to the side with more total flashes) were rewarded with points, which accumulated over trials. **E.** A treasure map informed players about game progress through 10 levels (indicated by “X” and a skull). **F.** The choice rule was not given verbally; participants learned through trial-and-error. Based on how rodents were trained in previous versions of the pulse-based task, we used the first 10 trials for training (level 1). The numbers of flashes on the two sides in the training trials were 10 vs 0 or 9 vs 1. For the remaining 190 trials (levels 2-10), the flash ratio between the two sides averaged at 70:30. There were at most 10 flashes per side per trial. Flash numbers were generated with a Poisson process.

To improve data collection, accessibility and engagement, we implemented several key design features. First, the game incorporated a feedback-based training pipeline to enable participants to learn the rules of the game without verbal instructions. Second, the game was published online, allowing participants to play from home, with an experimenter available over video call. Third, we developed an engaging graphical environment and storyline. Fourth, the game was playable via mouse or touch screen. Together, these innovations established a high throughput system that can help advance understanding of sensory perceptual deficits across the range of phenotypic heterogeneity of ASD (**Fig. 1C**).

From the perspective of the player, the game presented a quest to collect gems (**Fig. 1D**). Participants initiated each trial by clicking on an avatar at the center of the screen (initiation), after which the flashes, in the shape of gems, appeared bilaterally (cue period). After the cue period, the participant made a left/right choice and clicked on the chosen side. Correct choices were rewarded with points, which accumulated over trials. Participants were informed of game progress through a treasure map with 10 “levels” (**Fig. 1E**). To help participants learn the game mechanics and the response rule (choose the side with more flashes), we provided a learning curriculum in the form of training trials (game level 1, trials 1-10). The training trials presented the simplest stimuli, with most flashes appearing on one side (10 vs 0 flashes, or 9 vs 1 flashes). After the training (game levels 2-10), stimuli on each trial were determined by a stochastic process in which the probability of a flash (p) at each timepoint was 0.7 for the correct side and 0.3 for the incorrect side (**Fig. 1F**). Criteria for successful play were defined as completing all 200 trials and achieving an average accuracy of >=61.6% post training (i.e. trials 11-200).

In addition to game data, we collected four ASD-relevant surveys, filled out by the parents or guardians online: Social Responsiveness Scale (SRS-2; Constantino & Gruber 2012), Vineland Adaptive Behavior Scale (VABS-3; Sparrow et al. 2016), Adult Adolescent Sensory Profile (AASP; Brown & Dunn 2002), and Behavioral Inflexibility Scale (BIS; Lecavalier et al. 2020). SRS-2 was used as an index for ASD symptom severity. VABS-3 was treated as an index of level of adaptive functioning, and we used a threshold 75 to distinguish participants functioning at a higher level (VABS-3 >= 75) from those with significantly lower adaptive behavior including limited communication, social and daily living skills, which correlate with level of intellectual disability (VABS-3 composite score < 75).

### Individuals across the ASD spectrum learned the game successfully

Game players were recruited by emails to eligible families across a combination of databases that included SPARK, Simons Searchlight, the Phelan McDermid Syndrome data hub, and the Boston University Center for Autism Research Excellence registry. In all *N* = 308 adolescents (11 to 17 years, 198 male, 90 female, 20 unidentified), including *N* = 212 with a diagnosis of ASD, and *N* = 96 TD siblings played the game (**Supplementary** Fig. 1). To further characterize these participants, we collected VABS-3 scores from a subset of individuals (N=85 TD and N=195 ASD). The ASD sample spanned the spectrum, as reflected in SRS-2 and VABS-3 scores (**Fig. 2A–B; Supplementary** Fig. 2) and included a small number of individuals with Phelan McDermid Syndrome (PMS; N=3), and 16p deletion syndrome (N=3). Over 90% of TD (*N* = 79/85) and ASD (*N* = 87/96) participants with higher adaptive behavior scores (VABS-3 > 75), and more than 75% of ASD participants (*N* = 75/99) with lower adaptive behavior scores (VABS-3 < 75) were above criteria for success (**Fig. 2B**). This latter group included some whose VABS-3 scores were below 50, in the profound autism range (Lord et al., 2022). Both groups of ASD participants and TD participants who met criteria appeared to learn the rules of the game rapidly, within a few trials, and there were no significant differences in performance among the three groups during training (trials 1-10; **Fig. 2D**).

**Figure 2.**
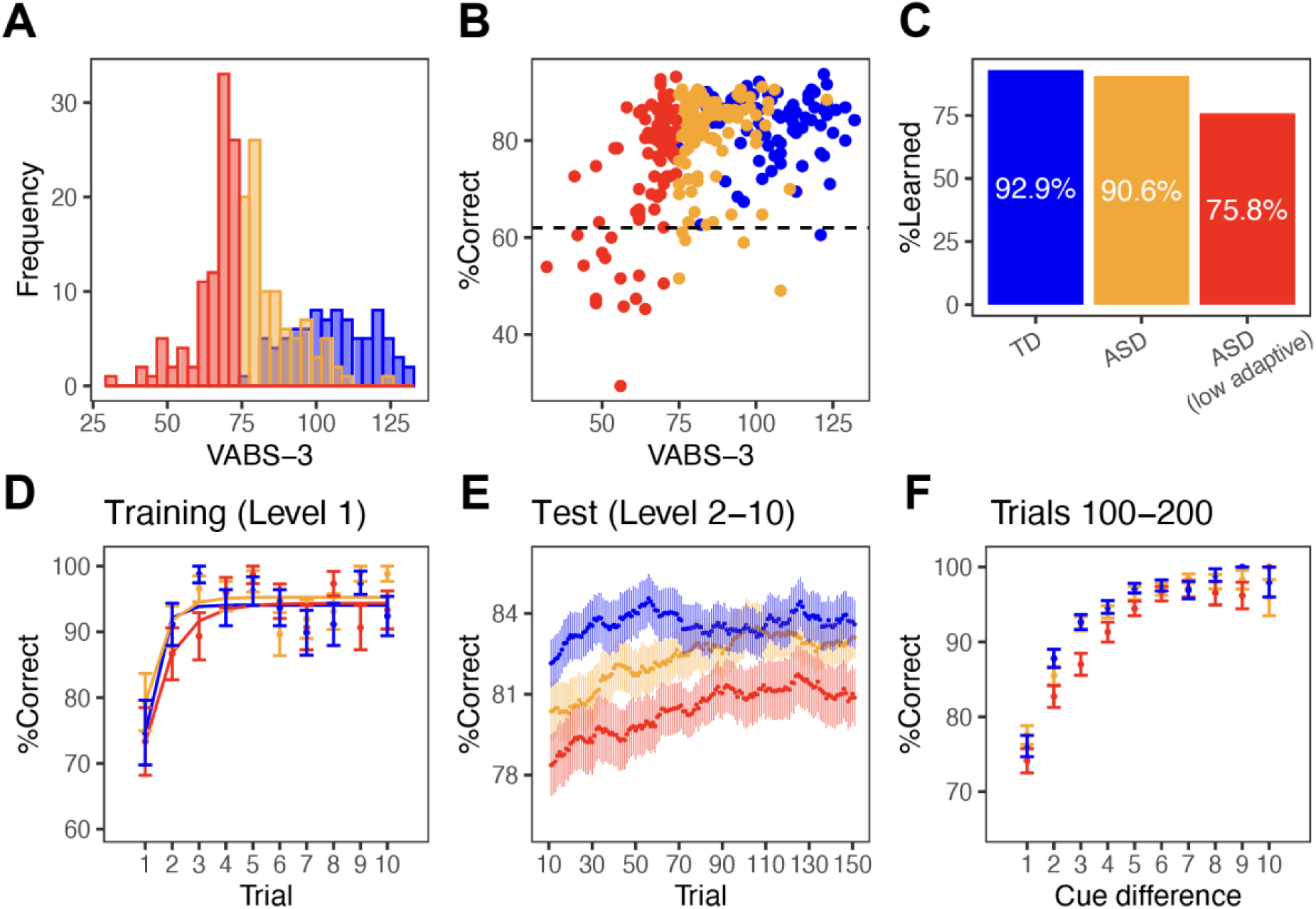
Participants across the ASD spectrum learned the game. **A.** We recruited 218 adolescents with ASD and 99 typically developing (TD) siblings. ASD varied in symptom severity, reflected in their VABS-3 score (absolute composite). We categorized participants into three groups: TD, ASD (with VABS-3 standard scores 75 or higher), and ASD low adaptive (with VABS- 3 lower than 75). **B.** The majority of participants in each group achieved an accuracy rate of 0.62 (upper limit of 95% binomial CI for chance performance) or above. **C.** Most participants passed task criteria (complete all 200 trials and average accuracy over 62%). **D.** Participants who passed criteria performed similarly across groups during training (level 1). **E.** The three groups showed differences in learning rate and overall performance during test trials (levels 2 to 10). ASD participants showed slower learning compared to TD participants, and ASD participants with low adaptive behavior scores differed even more. Plot shows rolling averages of accuracy over 50- trial bins. **F.** Psychometric function plots the proportion of correct choices as a function of the flash or cue (absolute) difference. Error bars represent SEM.

After transition to the full stimulus set, participants showed abrupt declines in accuracy, followed by a gradual increase up to a plateau level (**Fig. 2E**). This decline was greater for ASD than for TD players, and ASD players, on average, achieved lower plateau performance after recovery as assessed by both accuracy (**Fig 2F**) and the slope of the psychometric function (**Supplementary** Fig. 3). Furthermore, these deficits were consistent within participants across retest in a subset of participants (**Supplementary** Fig. 4). Examination of learning curve parameters after transition to the full stimulus set revealed significantly lower initial accuracy, learning rate, and plateau performance between the ASD and TD groups (KS test; p < 2.2e-16; see methods). These data indicate that participants across the ASD spectrum were able to successfully learn and play the game, but that, on average, ASD participants exhibited slower learning and slightly lower accuracy compared to TD participants.

### Individuals with ASD showed deficits in evidence accumulation

We next leveraged the varied and precisely known number and timing of sensory pulses on each trial to better understand participants’ perceptual decisions (**Fig. 3**). First, we asked how the timing of sensory evidence influenced decisions across the two groups of ASD and the TD participants. To capture the influence of sensory evidence at different times, we fit a logistic regression to pooled choice data from each group. In this model, predictors were binary variables indicating presence or absence of a flash in an individual time (250 ms bins), separately for left and right sides (**Fig. 3A**). After fitting, the model revealed similar integration time courses for each group. Integration was largely stable, with a slight recency and primacy bias that was consistent across groups. However, both groups of ASD participants had overall lower weights compared to TD (**Fig 3B**). There was a significant main effect of group [*F*(2,24) = 13.94, *p* < 0.001]; post hoc pairwise comparisons suggested significant differences between TD and lower adaptive functioning ASD participants (*p* < 0.001), as well as between the two ASD groups (*p* < 0.01). These results also held when early, middle, and late flashes were grouped together into three bins (**Fig. 3C**). These results suggest that, on average, all groups solved the game using a similar accumulation strategy in which pulses throughout the trial contributed to decisions. For the ASD groups, the lower regression weights could suggest other factors disrupting the integration process besides the timing of the flashes, not yet accounted for.

**Figure 3.**
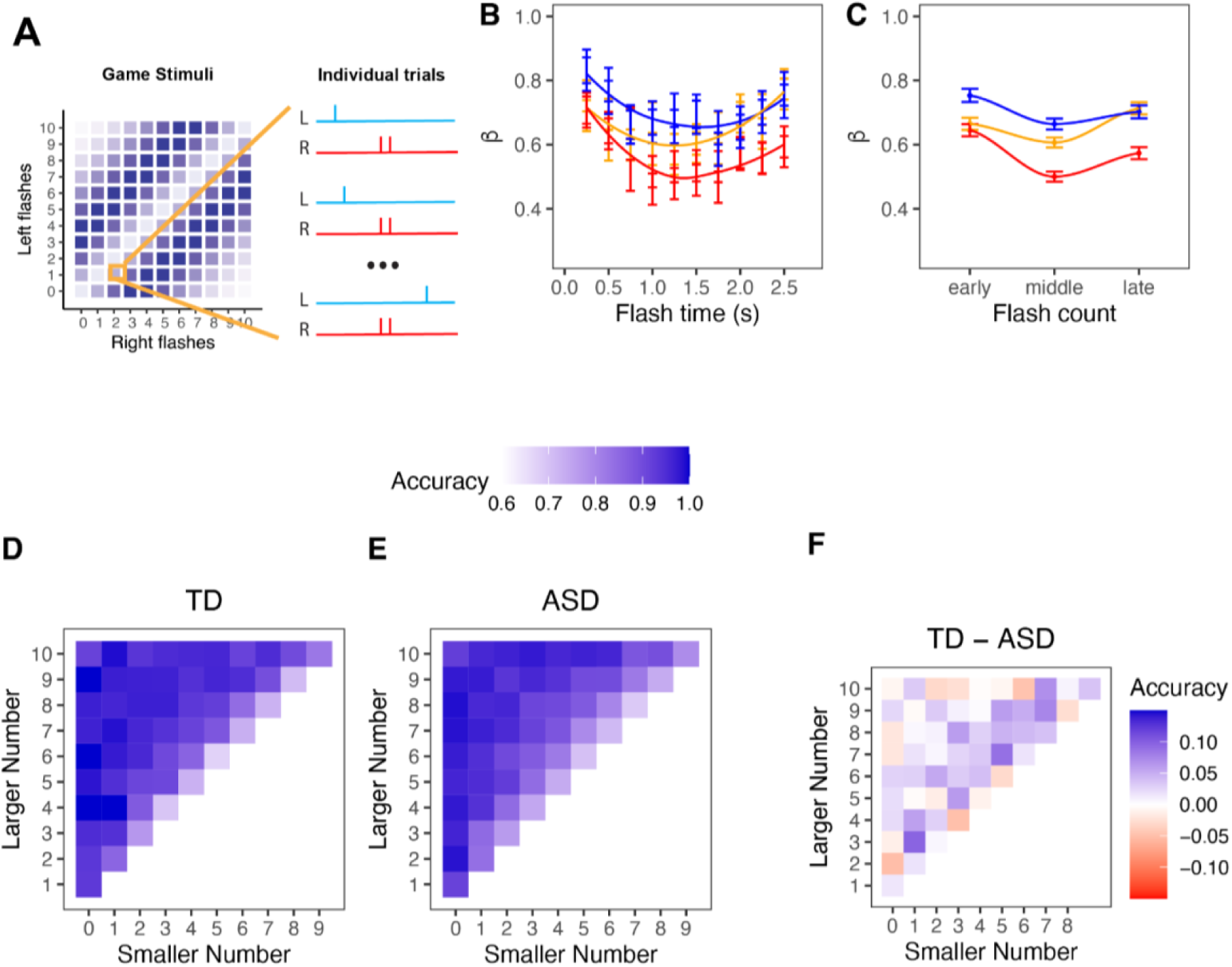
ASD participants showed deficits in evidence accumulation. **A.** The pulse-based paradigm incorporated high-dimensional stimuli; on each trial, sensory information was presented at different time points; each flash lasted 50 ms and flashes occurred in time bins of 250 ms. Trial-by-trial stimulus details facilitate characterization of participants’ perceptual choice behavior. **(B).** A logistic regression model estimated the weight (beta) of each flash (bin) on behavioral choice for TD (blue), ASD (gold) and ASD low adaptive (red) participants. **(C)** Logistic regression model for behavioral choice grouping early, middle and late flashes together. Models used data pooled across participants, model weights were collapsed across left and right sides, error bars are SEM of the average weight (for left and right), calculated using error propagation method. **D–F.** Two-dimensional psychometric function plot showing choice accuracy as a function of the number of flashes on each side, separately for TD **(D)**, ASD **(E)**, and the difference between TD and ASD **(F)**, data were pooled across participants in each group.

Next we sought to understand how the number of pulses influenced the decisions of ASD and TD participants. We evaluated three-dimensional psychometric functions that took into account how accuracy changed with the total number of flashes presented on each side (**Fig. 3D-E**). Comparing the performance of TD vs ASD (**Fig. 3F**), we found that ASD participants showed lower accuracy both as a function of decreasing difference between the number of left and right flashes and increasing total number of flashes. Together these data from both the ASD and TD groups are consistent with a noisy integration process in which noise depends on the number of accumulated sensory stimuli.

### Greater integration noise explains deficits in evidence accumulation in ASD

To better characterize the nature of noise in the integration process across ASD and TD players, we analyzed behavioral choices using a signal detection theory framework (Scott et al. 2015). In this framework, the perception of each sequence of flashes was modeled as a real number drawn from a gaussian distribution, reflecting uncertainty in the participant’s estimate of the number of flashes (**Fig. 4A–C)**. The mean of the gaussian was set equal to the true number of flashes and the variance was fit to the data. Greater variance corresponds to greater subjective uncertainty. We first evaluated two models, *linear variance* and *scalar variability*, which parameterize uncertainty differently. In the linear variance model, variance is proportional to the number of flashes (**Supplementary** Fig. 4). In the scalar variability model, inspired by Weber-Fechner scaling, variance is proportional to the square of the number of flashes. Model comparison revealed that the scalar variability model provided a better fit for both ASD and TD participants (**Supplementary** Fig. 4). This result, which matches previous findings in rodents (Scott et al. 2015) and human adults (Do et al. 2023), indicated that perceptual errors grew in a nonlinear fashion, consistent with noise in the integration process rather than in bottom-up sensory signals.

**Figure 4.**
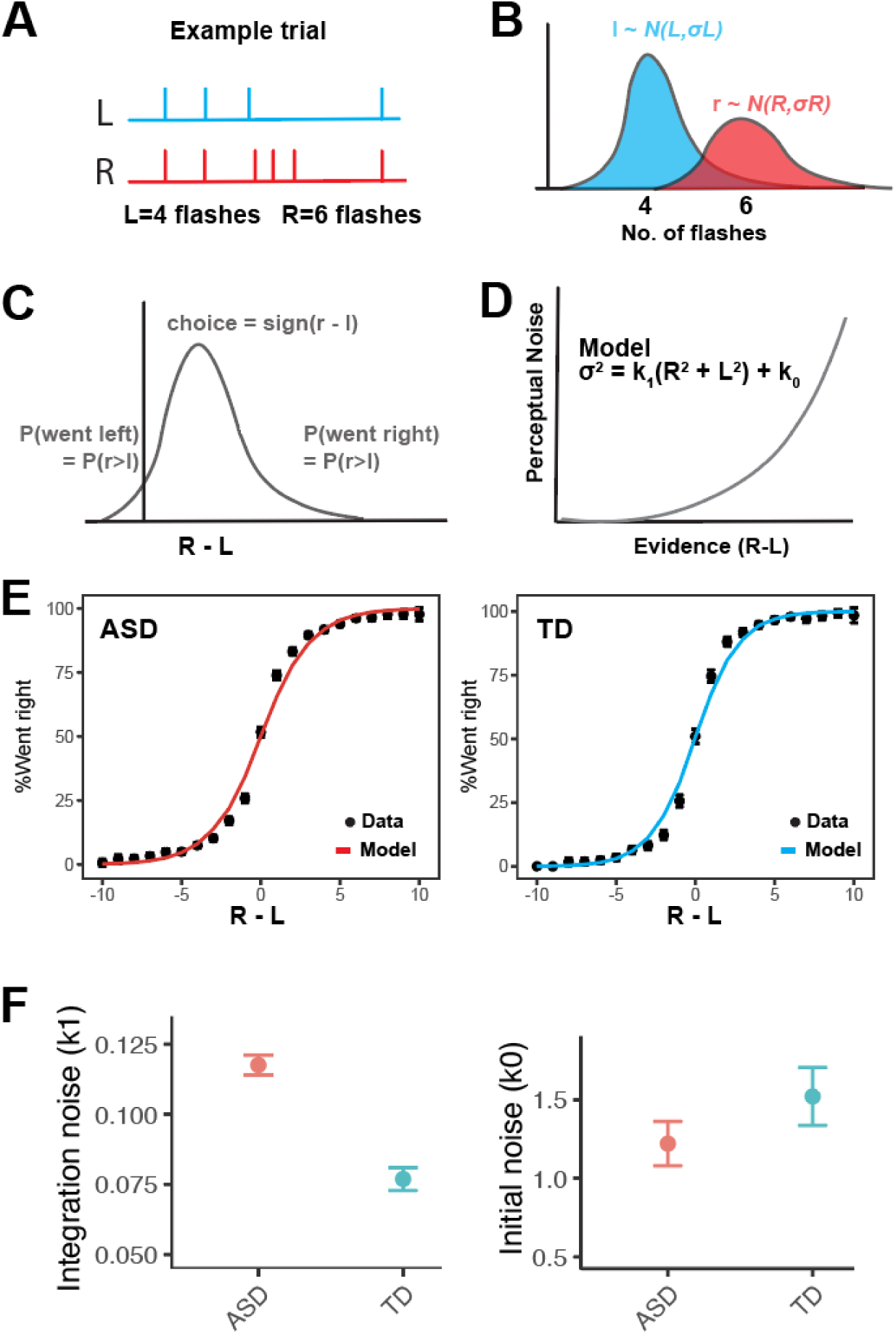
ASD participants experienced greater integration noise. **A.** Schematic of an example trial with 4 flashes on the left and 6 flashes on the right. **B–C.** A signal detection theory (SDT) model estimated flashes from gaussian random variables, with the mean reflecting evidence strength and variance reflecting internal noise. **D.** The model assumed scalar noise, associated with a free parameter (𝑘1). Other free parameters were side-bias (𝑏𝑠), win-stay (𝑏1) and lose-switch (𝑏2) biases. Model was fit to pooled data from post-training trials, for participants who met task criteria. **E.** Model fits (solid line) versus original data (error bars) for the ASD and TD groups. **F.** Model parameters for ASD and TD revealed greater integration noise in ASD. All error bars are SEM.

Next, we compared the magnitude of the integration noise across ASD and TD cohorts using the scalar variability model **(Fig. 4E**). This model included free parameters for integration noise (𝑘1) along with terms to capture side-bias (𝑏𝑠), a win-stay strategy (𝑏1), lose-switch strategy (𝑏2) and noise from all other sources (𝑘0). Parameter fits to pooled data revealed higher integration noise (𝑘1) (**Fig. 4F**) and win-stay (𝑏1) bias for ASD compared with TD, but no significant group differences in win-stay strategy (𝑏1), lose-switch strategy (𝑏2) or other sources of noise (k0). These results are consistent with the idea that greater noise in the integration process drives a deficit in perceptual decisions and evidence accumulation in ASD.

### Hierarchical Bayesian modeling reveals individual differences in evidence accumulation

Next, we asked how integration noise varied among individuals across the ASD spectrum. To address this question, we combined our signal detection theory approach with hierarchical Bayesian modeling (HBM-SDT) (**Fig. 5A–B**). By leveraging group-level and individual-level data, this approach can yield reliable noise parameter estimates for individual participants with small numbers of trials (Do et al., 2023). After fitting, the HBM-SDT model suggested wide variation in the parameter estimates for the ASD group (**Fig. 5C**). The integration noise parameter (𝑘1) was significantly greater for ASD than TD [Welch two sample test, 𝑡(252.46) = 2.46, *p* = 0.015]. Also, there was greater magnitude in side-bias [𝑏𝑠; 𝑡(227.34) = 2.03, *p* = 0.04], win-stay bias [𝑏1;𝑡(233.83) = 3.03, *p* = 0.003] and lose-switch bias [𝑏2; 𝑡(231.18) = 2.75, *p* = 0.006] for ASD compared with TD. Further, the fitted noise parameter from the HBM-SDT model showed good correspondence with the psychometric slope of individual participants, estimated with a generalized linear model (GLM; **Supplementary** Fig. 5). Finally, metrics were consistent across retest (**Supplementary** Fig. 6). Thus, the HBM-SDT model further supported increased integration noise in ASD and was able to capture the individual differences among ASD participants.

**Figure 5.**
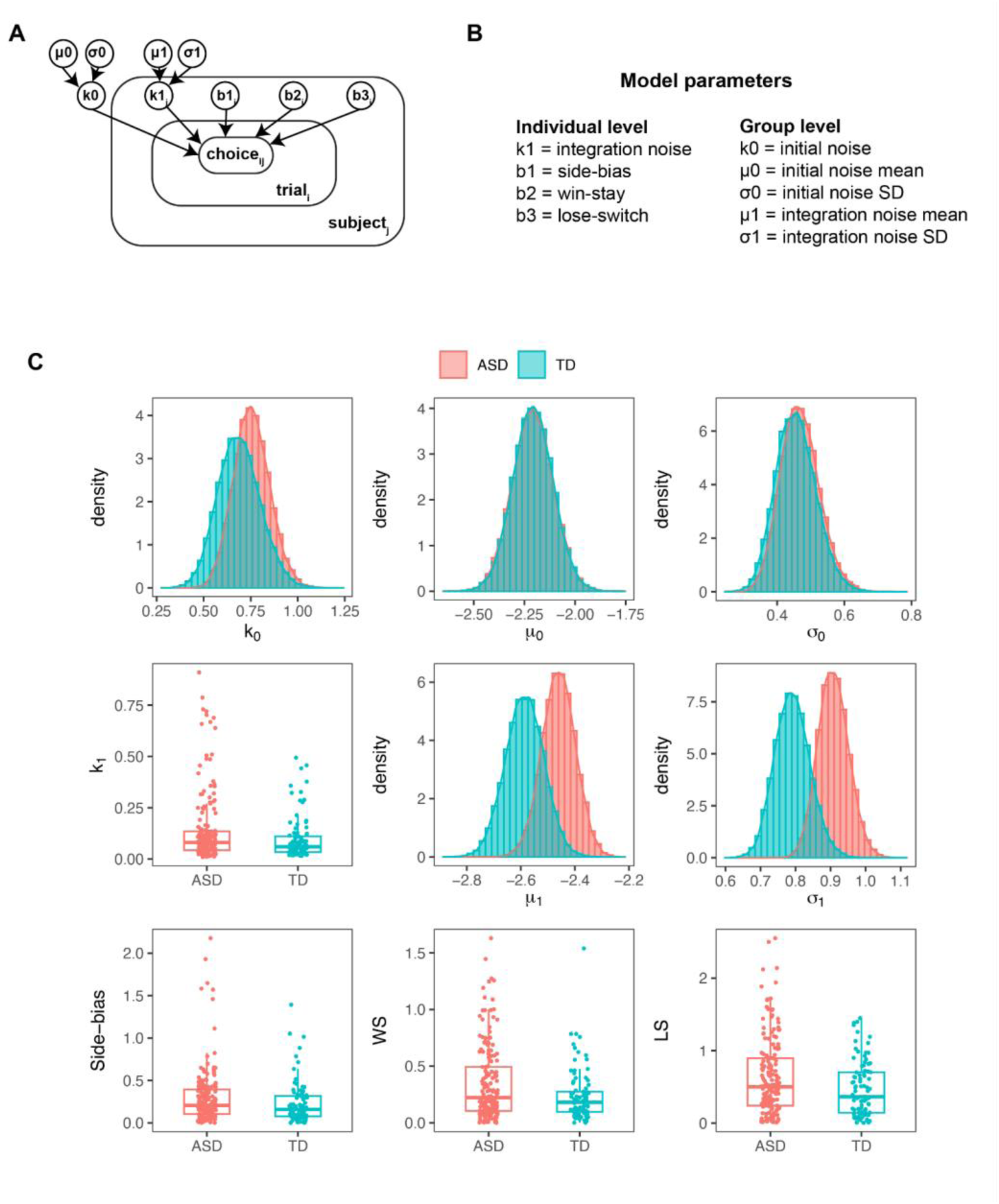
A hierarchical Bayesian model estimated individual differences in evidence accumulation. **A.** To fit the SDT model to individual participant data, we used a hierarchical Bayesian approach. **B.** The HBM-SDT model included different parameters, some estimated at the population level (histograms, indicating posterior probabilities), and some at the level of individual participants (box and whisker plots). The models were fit separately for ASD and TD. **C.** Fitted model parameter values suggested overall greater variability for ASD; significant group differences were also noted.

### Game-based measures correlated with survey scores

We next asked whether game metrics correlated with ASD symptom severity across individuals, as measured by survey scores. Thus, we analyzed the correlations between scores from SRS-2, VABS-3, AASP and BIS and the following game-based measures evaluated for individual participants: average accuracy, average intertrial interval (ITI, i.e., time taken to initiate trials), psychometric slope, integration noise, side-bias, win-stay bias, and lose-switch bias. Game measures correlated most strongly with VABS-3, followed by SRS-2 (**Fig. 6A–F**). Accuracy and psychometric slope also correlated with BIS scores (**Fig. 6G**). Interestingly, we failed to find significant correlations between game metrics and sensory profile scores (AASP). Overall, these correlations suggest that the game is useful for obtaining objective, quantitative parameters that correlate with traditional survey-based symptom severity measures related to both ASD (i.e. SRS- 2 and BIS) and level of adaptive functioning (i.e. VABS-3).

**Figure 6.**
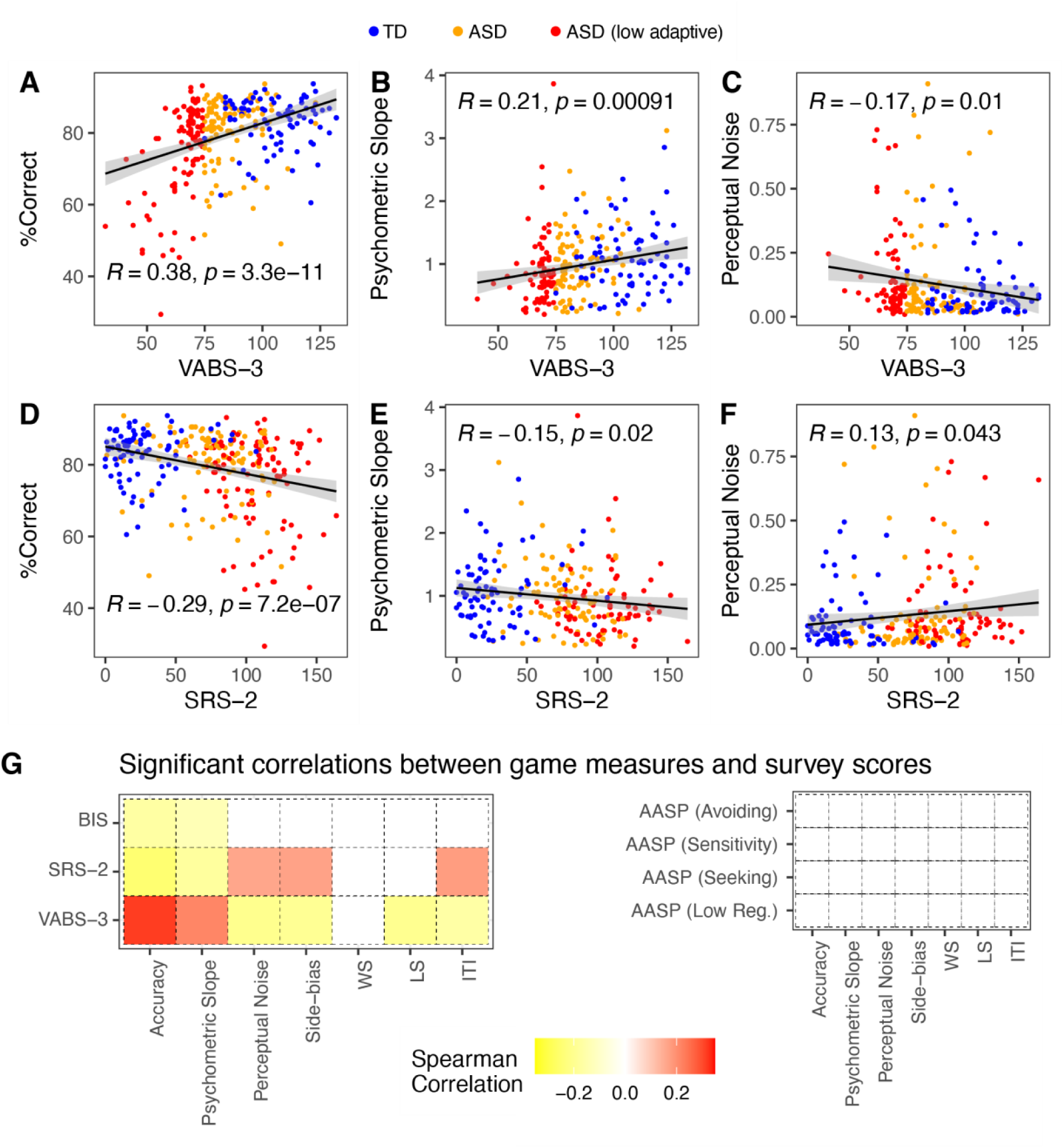
Game play correlated with symptom severity. A–F. Correlation between game statistics and ASD symptoms from caregiver surveys. Each colored dot represents a TD (blue), ASD (orange) or low-adaptive ASD (red) individuals. Black line indicates the slope. **G.** Correlation matrix between all computed game statistics and caregiver surveys. Color indicates the direction and strength of the correlation. White indicates no significant correlation (p>0.05). No significant correlations were observed between game statistics and sensory profile scores across individuals (p>0.05).

### Game measures explain additional source of variation between ASD and TD

Next, we wondered if the game-based measures could explain additional sources of variation among participants, beyond what is captured by the surveys. We conducted a principal component analysis (*PCA*) on the combined features (game and survey) from all participants with complete survey data (*N* = 243; 160 𝐴𝑆𝐷). The first two *PCs* explained 32% and 22% of the variation in the data respectively (**Fig. 7A**) and the ASD and TD groups were well separated in the *PC* space (**Fig. 7B**). Survey measures loaded heavily onto *PC1*, whereas game measures loaded heavily onto *PC2 (***Fig. 7C**). Furthermore, feature scores for *PC1* were significantly different for ASD and TD [Welch two sample test, 𝑡(158.95) = −4.10, *p* < 0.001]; this was also true for *PC2* [𝑡(194.25) = 2.29, *p* = 0.02]. This suggests that game-based measures revealed additional sources of variance in behavior that were not fully captured by the conventional surveys used in this study.

**Figure 7.**
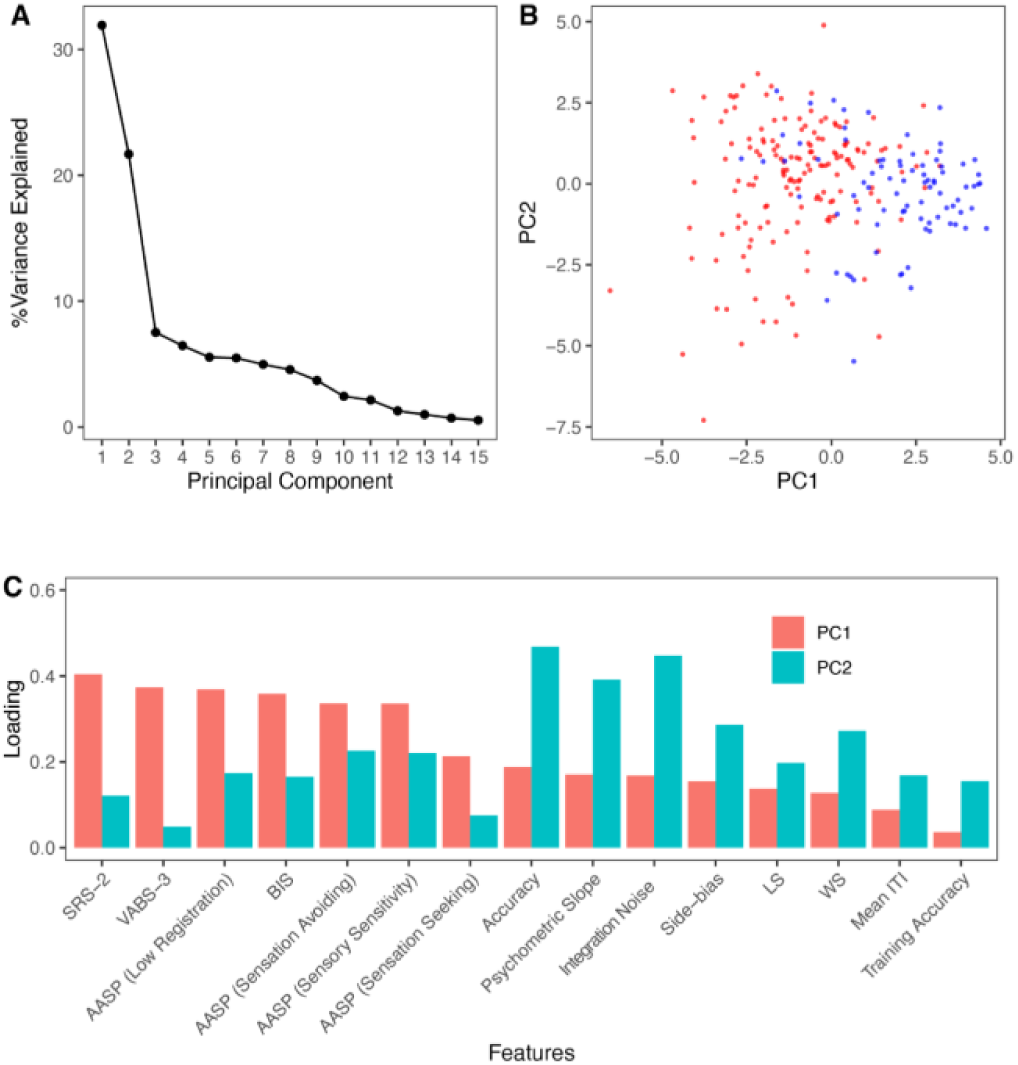
Game measures describe additional independent variance between ASD and TD. **A.** We conducted principal component analysis (PCA) on the combined (game and survey) features. Principal component (PC) 1 and PC2 explained most variance in the data. **B.** ASD individuals (red dots) and TD individuals (blue dots) were well separated in the space of the first two PCs. **C.** The survey measures loaded heavily onto PC1 (red) while the game measures loaded heavily onto PC2 (green). The PCA calculation did not use group information, this information was used to evaluate group differences in PC scores and for visualization.

#### Using artificial learning agents to model ASD traits

To better understand the relationship between integration noise and slower learning in our task, we modeled the behavior of TD and ASD populations using artificial neural network-based agents (**Fig. 8**). Like the human participants, the goal of these agents was to learn to associate sensory cues with the correct binary choice using feedback .

**Figure 8.**
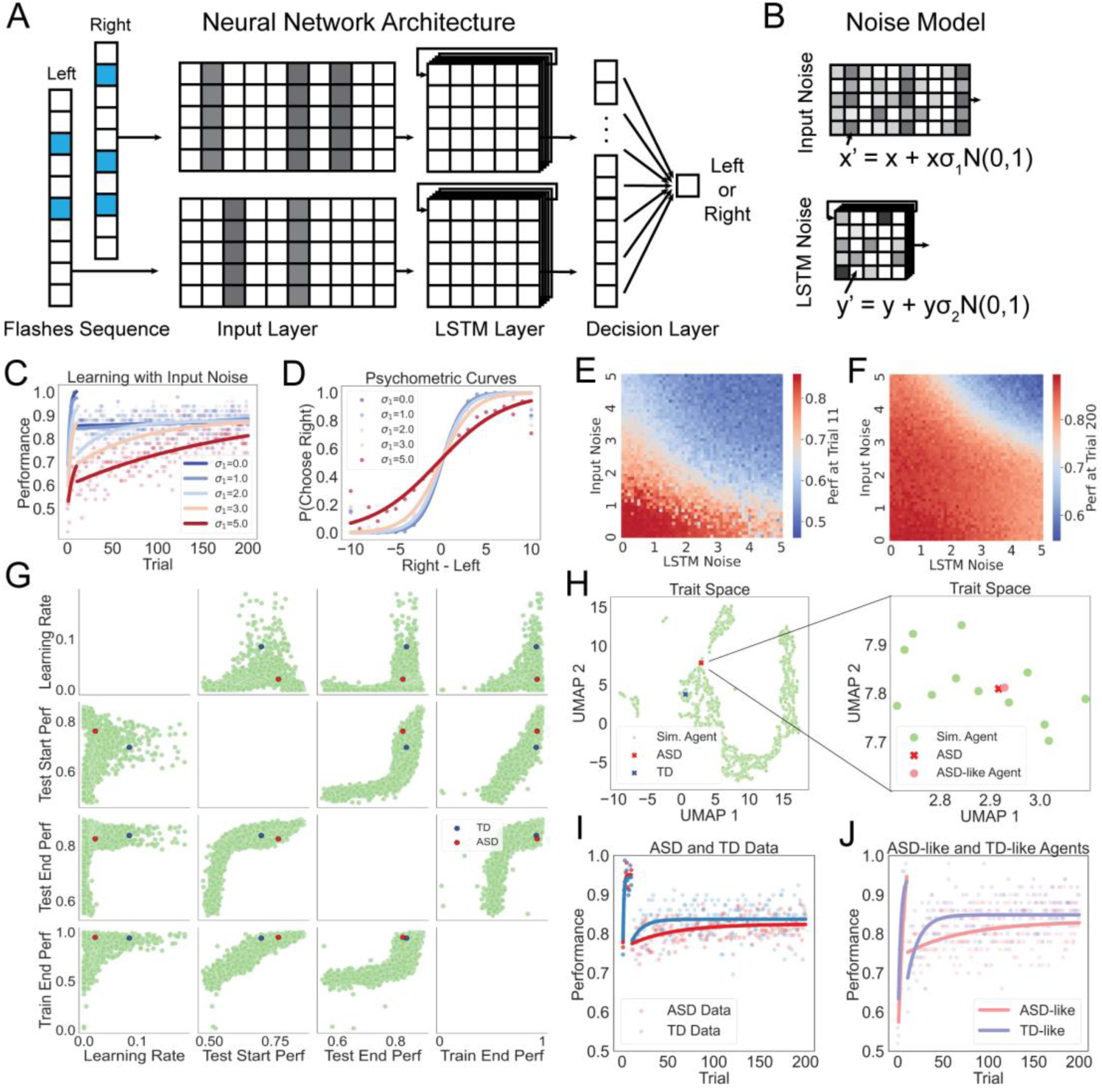
Using artificial learning agents to model ASD traits. (A) Neural network architecture for artificial learning agents. (B) Gaussian noise was injected into the input layer and LSTM layer. (C) Learning curve with varying input noise levels (σ_1_=0,1,2,3,5) across 200 trials, showing that artificial agents could be trained to perform the task using the same curriculum as human participants, and input noise altered both the rate of learning and plateau performance. (D) Psychometric curves fitted to models with different noise levels. (E - F) Heatmaps showing accuracy at trial 11 (E) and trial 200 (F) for agents with varying levels of input and LSTM noise. (G) Scatter plot illustrating the relationship between different performance metrics (traits) of simulated agents (green dots), with average traits for ASD (red) and TD (blue) participants. (H) *Left:* UMAP projection of simulated agents in trait space, with ASD (red) and TD (blue) plotted within this space. *Right:* Zoomed-in visualization of the ASD-like agent, identified via Euclidean distance search in trait space. (I) Performance over 200 trials for ASD (red) and TD (blue), with fitted learning curves. (J) Performance over 200 trials for ASD-like and TD-like agents, identified from the closest simulated matches, with fitted learning curves for comparison.

In these networks, the flash sequences for the left and right sides are processed through an input layer, followed by an LSTM layer and a fully connected decision layer that determines the choice (**Fig. 8A**). All layers are fully connected, allowing the model to integrate temporal and spatial information within and across trials. Initially, the parameters (weights between layers) are randomly assigned, and the agents must learn to perform the task using the same curriculum as the humans—10 training trials (level 1) followed by 190 test trials (levels 2–10). Learning occurs through feedback: at the end of each trial, the agent generates a decision, receives a reward signal (0 or 1), and updates its parameters using backpropagation with gradient descent.

To assess how integration noise affected the neural network’s performance, we injected Gaussian noise into early stages of processing: input noise was added to the input layer, affecting sensory representations, while LSTM noise was injected into the recurrent layer, influencing temporal integration (**Fig. 8B**). Noise strength was controlled by the standard deviations, σ₁ and σ₂, respectively.

In the absence of noise (σ₁ = σ₂ = 0), the model rapidly learned the task, reaching 85.53% accuracy at trial 11 (**Fig. 8C**). However, when noise was introduced, the learning rate and performance of the model changed (**Fig. 8C–D**). These effects followed a gradient, with increasing noise systematically altering both learning and integration (**Fig. 8E–F**).

We simulated a set of artificial agents with varying noise levels and mapped their positions in trait space, a five-dimensional space defined by behavioral performance on the task (i.e. accuracy at trials 10, 11, 200 and learning rate, **Fig. 8G**). Next, we identified the agents most similar to ASD and TD humans using a nearest neighbor approach (**Fig. 8H**). These ASD-like and TD-like agents captured several features of their human counterparts, including slower learning and lower peak performance in ASD participants (**Fig. 8I–J**). These results indicate that adding noise to neural representations is sufficient to generate the deficits in learning observed in ASD individuals.

## Discussion

Our results reveal that deficits in perceptual integration are widespread across the ASD population and correlate with symptom severity in multiple domains, including social responsivity and adaptive functioning. Model-based analysis suggested that these deficits could have stemmed from increased noise in the integration process. Integration noise grows non-linearly with the total number of flashes, explaining why ASD participants performed similarly to TD individuals on simple training trials but exhibited greater deficits when processing more complex stimuli. Furthermore, we demonstrate that introducing integration-like noise into artificial neural networks induces ASD-like traits in these agents, including slower learning and errors in perceptual judgments.

Our modeling efforts do not directly implicate a specific circuit origin for the increase in integration noise; however, previous studies have reported that ASD individuals display higher trial-to-trial variability in cortical responses to sensory stimuli than non-autistic individuals (Milne 2011; Dinstein et al. 2012; Weinger et al. 2014). One potential source for this increased variability is a change in the excitatory-to-inhibitory ratio, leading to chaotic neural dynamics as excitation or gain increases (Rubenstein and Merzenich 2003; Sohal and Rubenstein 2019). Indeed, computational models indicate slow timescale changes in gain in neural networks can lead to multiplicative noise similar to that revealed by our analysis (Gorris et al. 2018). Taken together, these studies suggest a model in which ASD risk genes drive neural circuit level changes that increase variability in neural responses leading to atypical perceptual integration (Dinstein et al. 2015; Haigh 2018; Sohal and Rubenstein 2019).

Our findings were made possible by an online video game, inspired by animal tasks, designed to assess perceptual processing in a large, heterogeneous sample of adolescents with ASD. The utility and innovation of this approach are evident in several key findings. First, ASD participants, including those with low adaptive skills and profound autism, successfully learned and played the game. Second, the game effectively detected perceptual differences between ASD and TD individuals. Third, game-based measures correlated with symptom severity, as assessed by survey data.

Beyond the utility for perceptual integration, there are several promising lines of development for new online games for adolescents with ASD that incorporate the frameworks described here. First the game could be altered to present sensory evidence in the auditory domain. Indeed, the pulse- based task in rodents was originally presented as clicks instead of flashes, providing a useful point of departure (Brunton et al. 2013). Second, game mechanics could be altered to capture other metrics, such as response time. To this end, we have recently designed a free response version of the visual pulse-based task to capture choice and response times (Kane et al. 2024) and have synchronized this task across humans and rodents to assess decision thresholds across species (Chakravarty et al. 2024). Third, future work could back-translate other tasks, such as the two-step task (Miller et al. 2022) or context dependent flexible switching (Pagan et al. 2024), from rodents to humans to assess other aspects of perception and cognition in ASD. Finally, we also note that the online task offers the possibility of much larger-scale data collection than what we have demonstrated here. In the present study, participant enrollment was limited by the scheduling time of the project coordinator who administered the game over video call. However, future versions of the game can be operated online in an automated fashion, without the need for experimenter supervision (Do et al. 2023). Such automation could allow a high rate of data collection and might also increase participation from families who otherwise find it difficult to schedule appointments. However, such a solution would require validation and may necessitate additional functionality to confirm player identity and engagement.

Finally, we demonstrate the use of artificial learning agents that exhibit ASD-like traits, including slower learning and deficits in perceptual integration when sensory and integration layers are corrupted by noise. In the future, such agents could be refined to capture additional ASD-like traits by incorporating behavioral data from other games or by employing improved search strategies, such as directed function evolution (Castro et al., 2025). Once optimized, these agents could generate hypotheses for testing in human participants. An intriguing possibility is their potential use in designing more effective training curricula for adolescents with autism. In this study, we applied a simple heuristic for curriculum design—starting with a small, fixed number of easy trials. However, by leveraging artificial ASD-like agents, it may be possible to develop more efficient training procedures to enhance learning in individuals with ASD and other neurodevelopmental disorders.

## Acknowledgments

We would like to thank all game players and their families. We would like to thank Vanessa Torres- Lacarra for project management, recruitment and data collection. We would like to thank game testers and developers, particularly Sadie Johnson who designed the graphics for the game. We thank Phoebe Lesk for help with data collection. We would like to thank Pam Feliciano (SPARK), Mahfuza Sabiha (SPARK), Marti Luby (SPARK/Boston Children’s Hospital) and Kate Still (Phelan McDermid Syndrome Foundation) for help with participant recruitment. This work was supported by a SFARI award (#874568) to BBS, HTF and JTM.

## Author contributions

All authors contributed to the conception of the study. QD designed the video game. SC, QD and YL analyzed the data. HTF, JTM and BBS provided guidance on data analyses and interpretation of results. SC and BBS wrote the manuscript. All authors provided feedback on editing the manuscript.

## Declaration of interests

The authors declare no conflict of interest.

## Methods

### Participants

All procedures were approved by the institutional review board at Boston University (IRB #6236E). We recruited N = 218 adolescents (11 to 17 years, inclusive) with an ASD diagnosis, and N = 99 TD adolescent siblings. ASD and TD participants were recruited from the same four databases: 1) SPARK Research Match (Simons Powering Autism Research for Knowledge; N = 166), 2) CARE (Centre for Autism Research Excellence) at Boston University (N = 46), 3) Simons Searchlight (N = 3), and the Phelan-McDermid Foundation (PMSF) data hub (N = 3). ASD diagnoses was confirmed by SPARK, Searchlight, and PMSF. Alternatively, a community diagnosis was confirmed by the parent/caregiver. An initial screening was conducted with the parent/caregiver to determine if the participant may be able to play the game with minimal support. Informed consent/assent were obtained from both the participant and the parent/caregiver. The parent/caregiver also provided demographic information about the participant, and completed four surveys: 1) Social Responsiveness Scale (SRS-2), which was used to further support ASD diagnosis and provided a quantitative measure for ASD symptom severity, 2) Vineland Adaptive Behavior Scales (VABS-3), which measured overall adaptive skills, and was used as a proxy for IQ, 3) Adolescent/Adult Sensory Profile (AASP), which measured sensory symptoms, and 4) Behavioral Inflexibility Scale (BIS), which measured inflexibility or rigidness of behavior. All participants were required to have normal or corrected-to-normal vision. Participants with a seizure disorder were excluded from the study.

N = 6 ASD and N = 3 TD participants were excluded from further analysis due to either data error or a non-typical SRS-2 score; for ASD, SRS-2 < 20 was considered non-typical, for TD, SRS-2 > 120 was considered non-typical. Task success was determined by the following: completion of all 200 trials, and average accuracy of 61.6% or above for trials 11 through 200, this was based on the upper limit of the binomial 99% confidence interval for chance performance. N = 89 TD and N = 174 ASD participants achieved this criterion. Further, to separately evaluate the performance of more or less able ASD participants, we used their VABS-3 composite standard scores; ASD participants with a score of 75 or above were in the more able group, those below 75 were considered less able with low adaptive skills, including some with profound ASD. Note that N = 17 ASD participants had missing VABS-3 scores, N = 11 TD participants also had either missing or unusually (VABS-3 < 75) low scores. These participants were excluded from in which VABS-3 scores were used. To test whether participants who did not meet task criteria could do so on a second attempt we re-tested a subset of ASD individuals (N= 11 individuals; see **Supplementary** Fig. 1). To gauge the reliability of the game-based measures across multiple game plays we reached re-tested a subset of ASD and TD individuals who did achieve criterion on the first attempt (N= 35 individuals, 23 ASD; 12 TD; see **Supplementary** Fig. 6).

## Video game development

The online video game was inspired by a pulse-based evidence accumulation task for rodents (Scott et al. 2015; Scott et al. 2017; Scott et al. 2018) and modified from an earlier version used in humans (Do et al 2023). The game, playable on a computer as well as a tablet, has many visual details and a background narrative. It is a quest to collect gems. The task underlying the game includes streams of visual flashes. The flashes appear in 250 ms time bins. For a single trial, there could be up to 10 flashes on each side. Each trial was divided into three distinct states—trials began in the initial state, when the participant clicks or touches on the center of the screen. The trial then enters the cue state in which the participant is presented with the flash stimuli (in the shape of the gems). The cue state lasts for 2.5 to 3 s, after that the trial enters the decision state, during which the participant makes a left or right decision. Post decision, participants are provided with feedback in terms of point bonuses or no points for erroneous choices. Game progress is conveyed with a graphical treasure map, presented from time to time.

The game had a total of 200 trials, the first 10 trials were used for training: there were 10 vs 0 flashes or 9 vs 1 flashes on training trials. After training, participants were presented with the full stimulus set, where, on average, the flash ratio was 70:30. Flashes were generated using a Poisson process, there could be a flash on either side, or both, or none, for a particular time bin. The game is available to play online at http://eviaco.herokuapp.com/cave.

### Data collection

Data were collected online using a split screen video call (Zoom). At the beginning of a session, the experimenter confirmed the participants’ information, then forwarded the link to play the game. Participants were provided with a de-identified subject ID with which they used to log into the game. Parents/Caregivers received the following script:

“*First, we will go through the informed consent form. I will walk you through this and please feel free to ask any questions as they come up. If you would still like to participate once we’ve gone through the ICF, you can go ahead and sign whenever you are ready. While your teenager is playing the video game, we are going to have you fill out some questionnaires about your teenager’s behaviors. This should take around 30 minutes as they are all fairly short. If you have any questions about the various forms, feel free to ask me at any point. Once you finish the questionnaires, you are all set.”*

Participants were not provided with instructions on how to play the game, but were prompted on how to start the game using the following script:

“*Gathering Evidence for Optimizing Decisions is a study where you will be playing a video game. I’m going to explain the game a bit to you now! I*’*m not going to tell you very much about it, but I*’*ll show you how it works. Then I want you to see if you can figure out how to win the game! You have about 45 minutes to play.”*

Or, for adolescents with lower verbal abilities:

“*Gathering Evidence for Optimizing Decisions is a study where you will be playing a video game. I*’*m going to explain the game a bit to you now! When you click the screen, some flashes will come up and from there you have to decide what to do!”*

On the first few training trials, the experimenter provided verbal encouragement such as “Good job” for a correct response or “oops” for an incorrect response. The experimenter allowed the parent/caregiver to be present during game play, to provide additional support. Occasionally, the experimenter gave out the rule of the game (go with the side with more flashes) upon participant request. There were also a few instances where the parent/caregiver provided some guidance. Nonetheless, all these instances were logged in detail and accounted for post hoc in the analysis of the data. After completing the game, participants were thanked for their participation and rewarded with an Amazon gift card worth USD 25. If both siblings were invited to participate, sessions were held at different times/days.

The demographic information and survey responses were collected using REDCap (Research Electronic Data Capture) a secure online data management system (Harris et al. 2019). Responses made in the game, along with the game variables were collected using the MongoDB platform (https://www.mongodb.com/). Video and audio of the zoom sessions were also recorded and stored in a secured database. All data were de-identified.

### Data analysis

Data analyses were conducted using R Version 4.2.1; all Figures were generated using R package ggplot2 (Wickham, H., 2016).

***Analysis of survey scores.*** We followed published guidelines for calculating the scores for the four different surveys conducted in this study. SRS-2 is a 65-item questionnaire, each question is answered on a Likert scale, individual item scores are within the range of 0 to 3, some items (3, 7, 11, 12, 15, 17, 21, 22, 26, 32, 38, 40, 43, 45, 48, 52, 55) were reverse scored, missing item scores were replaced with the median values; items 52 and 55 had a median value of 2, whereas items 3, 5, 7, 9, 11, 12, 15, 17, 19, 21, 22, 25, 26, 28, 31, 38, 40, 43, 45, 48, and 56 has a median value of 1, all other items has a median value of 0. After these adjustments, the total score was calculated. SRS-2 total score served as an index for confirming ASD diagnosis, and thus, TD participants with extremely high SRS-2 scores (above 120) or ASD participants with very low scores (below 20) were considered as outliers. VABS-3 is a 120-item questionnaire, items are divided into three domains: communication, socialization and daily living skills. After calculating the raw totals for each domain, these were converted into standardized scores which corrects for age. Then, the standardized scores from three domains were used to calculate the Adaptive Behavior Composite (ABC) score, which was used in the analyses. AASP is a 60-item questionnaire, divided into four quadrants: low registration, sensory seeking, sensory avoiding, and sensory sensitivity. We used the raw totals for each quadrant. Finally, BIS is a 38-item questionnaire, the raw total score was used for BIS. Participants with missing SRS-2 scores were excluded from all analyses. Participants with missing VABS-3, AASP or BIS scores, were removed only from analyses that included those measures.

***Division of participant groups.*** Our main groups of interest were ASD and TD. As mentioned above, SRS-2 scores were used to support ASD diagnosis; a small number of participants with anomalous SRS-2 scores were excluded. We collected a heterogeneous sample of ASD participants, including participants with intellectual disability and who were minimally verbal. Thus, in analyses pertaining to group level differences, we divided the ASD group into two subgroups, based on their VABS-3 ABC scores, above or below 75 to capture more and less able participants. In our analyses pertaining to individual differences, these subgroups were dropped. Another division of participants was based on task success, participants who completed all 200 trials and achieved an average accuracy of 61.6% or above, were successful at the task, these participants were included in further analyses, average accuracy was the proportion of correct choices for trials 11 through 200.

***Average accuracy during training and post training.*** Motivated from staircase training curricula used in rodent experiments with the same task, here we included 10 training trials at the beginning, the first five trials had 10 vs 0 flash, and the next five had 9 vs 1 flashes. All participants experienced the same flash numbers for each training trial. To analyze training data, we averaged trial-wise accuracy across participants in each group. An exponential learning curve of the form 𝑃_𝑛_ = 𝑃_∞_ − (𝑃_∞_ − 𝑃_0_)𝑒^−ɑ𝑛^ was fit to the average accuracy using nonlinear least square estimation, where 𝑛 is the trial number, 𝑃_𝑛_ is the accuracy on trial 𝑛, 𝑃_∞_ and 𝑃_0_ are the asymptotic and initial accuracy, respectively.

For trials post training (trials 11 through 200), the flash numbers varied, though on average the flash probability on one side was *p* = 0.7, flash probability on the other side was 1 − *p*. To look at accuracy post training, we binned trials for each participant. A rolling average of accuracy over binned trials was used, with a bin size of 50 trials

### Bootstrapping and learning curve analysis

We fit an exponential learning curve model to participants’ trial-by-trial accuracy during the hard phase of the task (post-training phase) defined as:

y=A0 + (P−A0)*(1−e^(L*trial)) where:

where y is the accuracy at a given trial, A0 is the initial accuracy, P is the asymptotic (plateau) accuracy, and L is the learning rate. To estimate variability in these parameters at the group level, we performed bootstrap resampling by sampling participants with replacement within each group (ASD, TD), fitting the learning curve to each resampled dataset over 1,000 iterations.

Group differences in the resulting parameter distributions were assessed using the Kolmogorov–Smirnov (KS) test.

***Psychometric functions.*** We used psychometric functions to look at the relation between choice/accuracy and the sensory information, i.e., the flash numbers. Note that on each trial there could be at most 10 flashes per side. Trials with equal number of flashes on both sides were not rewarded. We fit a GLM to the psychometric data. The model estimated the probability of choosing the right side (𝜋) as a weighted sum of side-bias, flash difference (𝑅 − 𝐿), previous reward (𝑊𝑆), and previous omission (𝐿𝑆).

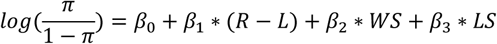

Here 𝛽 estimates the slope of the psychometric. 𝑊𝑆 was coded as: 𝑊𝑆 = 1 for a previous rewarded right choice, 𝑊𝑆 = −1 for a previous rewarded left choice, and 𝑊𝑆 = 0 otherwise. Similarly, 𝐿𝑆 was coded as: 𝐿𝑆 = 1 for a previous unrewarded right choice, 𝐿𝑆 = −1 for a previous unrewarded left choice, and 𝐿𝑆 = 0 otherwise. The GLM was fit to pooled data from each participant group, and for each trial bin of consideration.

***Flash weights on choice probability.*** To estimate the weight of each flash time bin on choice probability we used a binomial GLM, in which choice probability (𝜋) was modeled as:

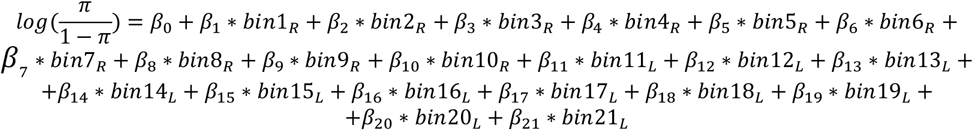

The bin parameters are binary, indicating whether there was a flash on a particular bin, each trial could be divided up into 10 time bins (250 ms each). The GLM was fit to pooled choice data from each participant group

***Signal detection theory model.*** We used a signal detection theory (SDT) approach to understand the influence of noise in participants’ choice decisions (also see Do et al., 2023). The model estimates choice probability (going to the right side over the left side) based on the probability of the perceived flash difference (𝑌) being greater than zero. Assume that the perceived number of flashes on the left [𝐿∼*N*(𝜂_𝐿_, 𝜎_𝐿_)] and right [𝑅∼*N*(𝜂_𝑅_, 𝜎_𝑅_)] sides can be treated as gaussian random variables. Then, the probability of going right is: 𝑃[𝑌 > 0] = 𝑃[(𝑅 − 𝐿) > 0]. 𝜂 and 𝜂 are the means, 𝜎_𝐿_ and 𝜎_𝑅_ are the SDs of the perceived flashes on the left and right sides. This makes 𝑌 also a gaussian variable, with mean (𝜂_𝑅_ − 𝜂_𝐿_), and SD √𝜎^2^_𝐿_ + 𝜎^2^_𝑅_, for when the noise is scalar. We added the win-stay, lose-switch and side-bias parameters to the difference in the mean (the evidence). The model was fit to pooled choice data from each group, using maximum likelihood estimate (MLE), with simulated annealing (SANN). We included all post-training trials from participants who met task criteria.

***Hierarchical Bayesian model.*** We used the RStan package (Stan Development Team, 2020) to fit a Hierarchical Bayesian Model (HBM) based on the SDT framework described above (see also, Do et al., 2023). Individual choices were modeled as a Bernoulli random variable, with probability 𝜋_𝑖𝑗_ – the probability of going to the right, as defined in the SDT framework. In the HBM, integration noise (𝑘1) was fit both at population- and individual subject level (Fig. 4–5). Initial noise (𝑘0) was fit at the population level only. The side-bias (𝑏1), win-stay (𝑏2) and lose-switch (𝑏3) parameters were added to the evidence (the mean), these were also estimated at the individual subject level.

𝑏1, 𝑏2, and 𝑏3 were assumed to be standard normal; 𝑘0 and 𝑘1 were assumed log normal with means and SDs shared across the population. HBM was fit separately to ASD and TD. We included all post-training trials from participants who met task criteria. Corrections for multiple comparisons were done through permutation tests.

### Neural Network Simulation

**Neural Network Architecture and Training.** To model learning in a sequential decision-making task, we implemented an artificial neural network with a dual-stream architecture, consisting of an input embedding layer, two independent LSTM layers, and a fully connected decision layer. The network processes sequences of visual stimuli and learns to make binary decisions based on these inputs.

1. **Input Representation.** The model processes two sequences of flash stimuli, one corresponding to the left side and one to the right. These sequences are first mapped into a continuous representation using two independent embedding layers.

1. ● The input size is 2, representing 2 binary flash sequences.
2. ● Each input is embedded in a 64-dimensional space.
3. ● The two embedding layers do not share weights, ensuring separate feature extraction for the two input streams.
2. **LSTM Layers for Temporal Processing.** After embedding, each sequence is passed through an independent Long Short-Term Memory (LSTM) layer:

1. ● Each LSTM layer has a hidden state dimension of 64.
2. ● The two LSTMs operate in parallel, processing the left and right input streams separately.
3. ● These LSTMs enable the model to integrate information across time steps and extract temporal patterns.
3. **Fully Connected Decision Layer.** The final decision is computed using a fully connected (linear) layer:

1. ● The outputs from both LSTMs are concatenated into a single feature vector.
2. ● Given that each LSTM processes 20 time steps (10 for each flash sequence), the concatenated input to this layer has a total size of 20×64+20×64=1280
3. ● The output dimension is 2, representing the probability of choosing the left or right option.
4. ● A softmax activation function is applied to obtain final decision probabilities.

The model is trained over 200 trials, with learning occurring after each trial. The training follows these steps:

1. Evaluation Phase:

1. ○ The model processes the input sequences and makes a binary decision (left or right).
2. ○ The decision is compared to the correct answer, and the model receives a score of 0 (incorrect) or 1 (correct).
2. Learning Phase (Backpropagation):

1. ○ Immediately after evaluating a trial, the model performs backpropagation based on the outcome of that specific trial.
2. ○ The network updates its weights accordingly before proceeding to the next trial.

This trial-by-trial learning setup allows the model to adapt incrementally, reflecting a continuous learning process similar to what happened during the video game that humans played.

**Noise Injection.** To examine the effects of stochasticity in the sensory representation on learning performance, Gaussian noise was introduced at multiple stages of the neural network. Specifically, noise was applied to both the input embedding layer and the LSTM layer to investigate how variability at different processing stages affects learning dynamics.

Noise was generated by sampling random values from a standard normal distribution N(0,1) where the mean is 0 and the standard deviation is 1. The noise was then scaled by a parameter σ, transforming the noise distribution to N(0,σ^2). This transformation ensures that the injected noise retains a mean of 0 while allowing precise control over its standard deviation through the parameter σ. The scaled noise was added directly to the input tensor x before being processed by the network.

To prevent the noise scaling process from interfering with gradient computations during backpropagation, the scaling factor σ was applied using a detached version of the input tensor. This ensures that the noise injection does not influence the optimization process, isolating its effects from the network’s learning dynamics.

Both input noise and LSTM noise were systematically varied across different conditions. Noise levels (σ) ranged from 0 to 5 in increments of 0.1, generating a wide spectrum of variability conditions for analysis.

**Two-Phase Learning Curve Fitting.** To quantify the relationship between trial number and accuracy, learning performance was modeled using an exponential learning curve function: y=A0 + (P−A0)*(1−e^(L*trial)) where:

● y is the accuracy at a given trial,
● A0 represents the initial accuracy,
● P is the plateau accuracy value,
● L is the learning rate, controlling the rate of improvement.

To account for differences in learning dynamics between early training and later generalization, the learning curve was fitted separately for two trial segments:

1. Trials 1–10 (Training Phase):

1. ○ These trials consisted of easier training conditions (9 vs. 1, 10 vs. 0), facilitating initial learning.
2. ○ The parameter A0 was fixed at 0.5, assuming chance-level performance at trial 1.
3. ○ The learning curve parameters extracted from this phase quantified the model’s adaptation to structured training.
2. Trials 11–200 (Testing Phase):

1. ○ Beginning at trial 11, the model was presented with the full stimulus set, requiring generalization beyond the initial training phase.
2. ○ A second learning curve was fitted to these trials to capture differences in long- term learning trajectories.

By fitting separate learning curves to the training and generalization phases, we obtained five parameters (A0_test, P_train, P_test, L_train, L_test) describing the learning behavior of each agent. These parameters were used as trait measures to characterize individual learning profiles.

**Simulation Design.** To systematically examine the effects of noise on learning, both input noise and LSTM noise were varied across a range of conditions:

● Noise levels were incremented from 0 to 5 in steps of 0.1,
● 50 agents were simulated for each noise condition,
● Learning curves were fitted to the averaged performance of each agent group.

This procedure generated a large-scale dataset of simulated agents, allowing detailed analysis of the impact of noise on learning performance.

In addition to the simulated agents, the same learning curve model was fitted to human behavioral data from typically developing (TD) and autism spectrum disorder (ASD) participants. This enabled direct comparison between human learning traits and those of the neural network agents.

**Nearest-Agent Matching Using cKDTree.** To identify simulated agents that best resembled the learning behavior of ASD and TD participants, we applied a nearest-neighbor search using the cKDTree algorithm from scipy.spatial.

### Matching Procedure

1. Each simulated agent was represented in a five-dimensional trait space , where the dimensions corresponded to the fitted learning parameters (A0_test, P_train, P_test, L_train, L_test).
2. The ASD and TD participants’ empirical data were projected into this trait space based on their fitted learning parameters.
3. Using Euclidean distance, the closest simulated agents to the ASD and TD data points were identified.
4. These nearest-matching agents were then analyzed to assess how well they captured ASD-like and TD-like learning dynamics.

This approach allowed us to systematically compare simulated and human learning behaviors, providing insights into how different noise levels influence learning trajectories.

### Dimensionality Reduction with UMAP

To visualize the high-dimensional trait space, we applied Uniform Manifold Approximation and Projection (UMAP), reducing the five-dimensional learning trait space to a two-dimensional (2D) representation.

UMAP Projection Procedure

● The five learning parameters extracted from the model were treated as feature vectors for each agent.
● UMAP was used to reduce these vectors to a lower-dimensional space while preserving their structural relationships.
● The projected 2D UMAP space was used to visualize the organization of ASD-like and TD-like agents.
● Nearest simulated agents were identified in this space by previously computing Euclidean distances between ASD, TD, and the generated agents in the original 5D space.

---

**Supplementary Figure 1.**
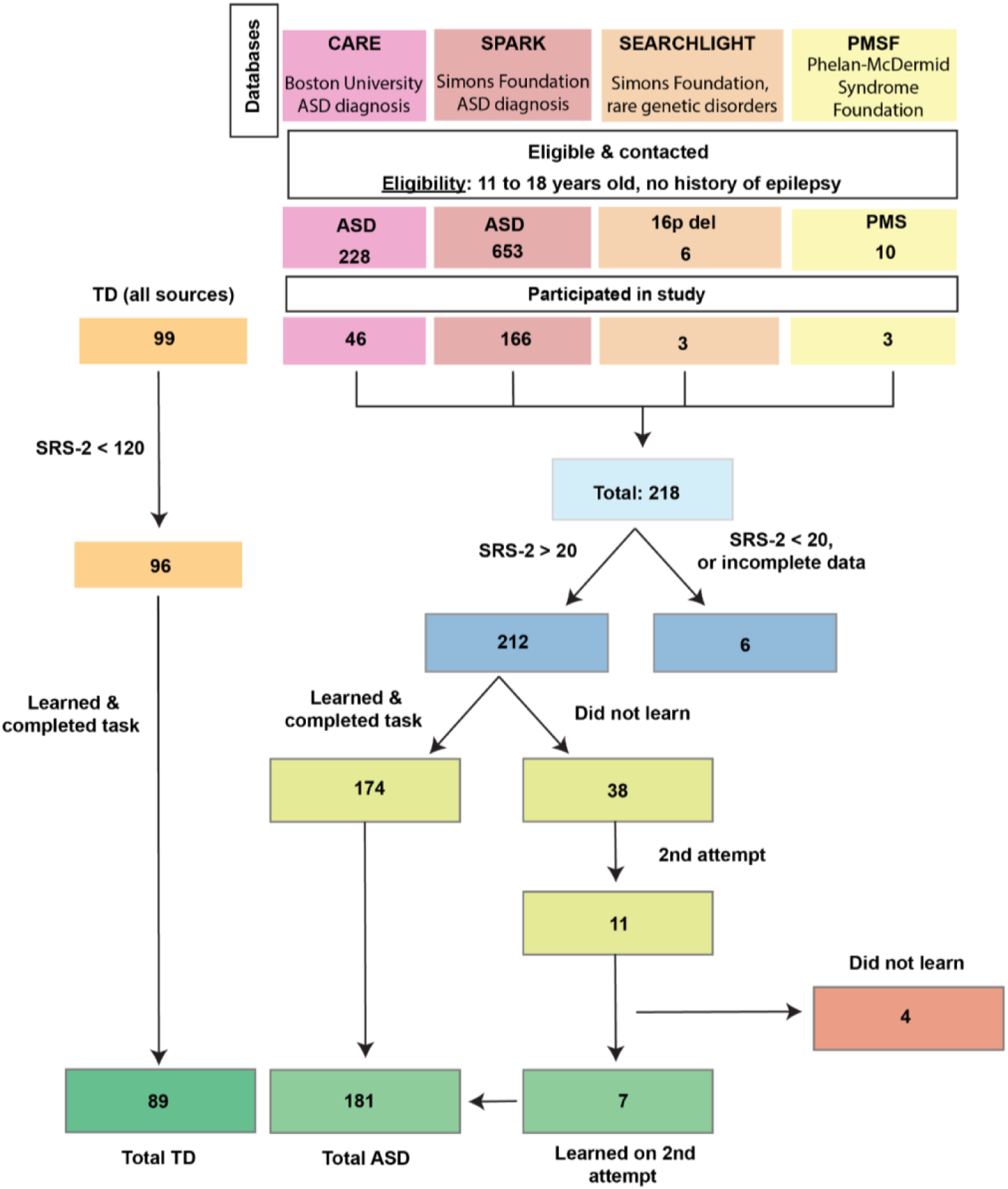
Recruitment. Participants were recruited from different databases, contacted through email advertisements. Eligibility was determined by age, ASD diagnosis and history of epilepsy. Those with missing game data were excluded. Participants with unusual SRS- 2 scores, defined as SRS-2>120 for TD (N=3) and SRS-2<20 for ASD (N=6) were also excluded. Criteria for successful game play and learning were determined by 1) completion of all (N = 200) trials, and 2) average accuracy of 61.6% or above post training. A subset of ASD participants who failed to reach above criteria were invited back for another session (N=11).

**Supplementary Figure 2.**
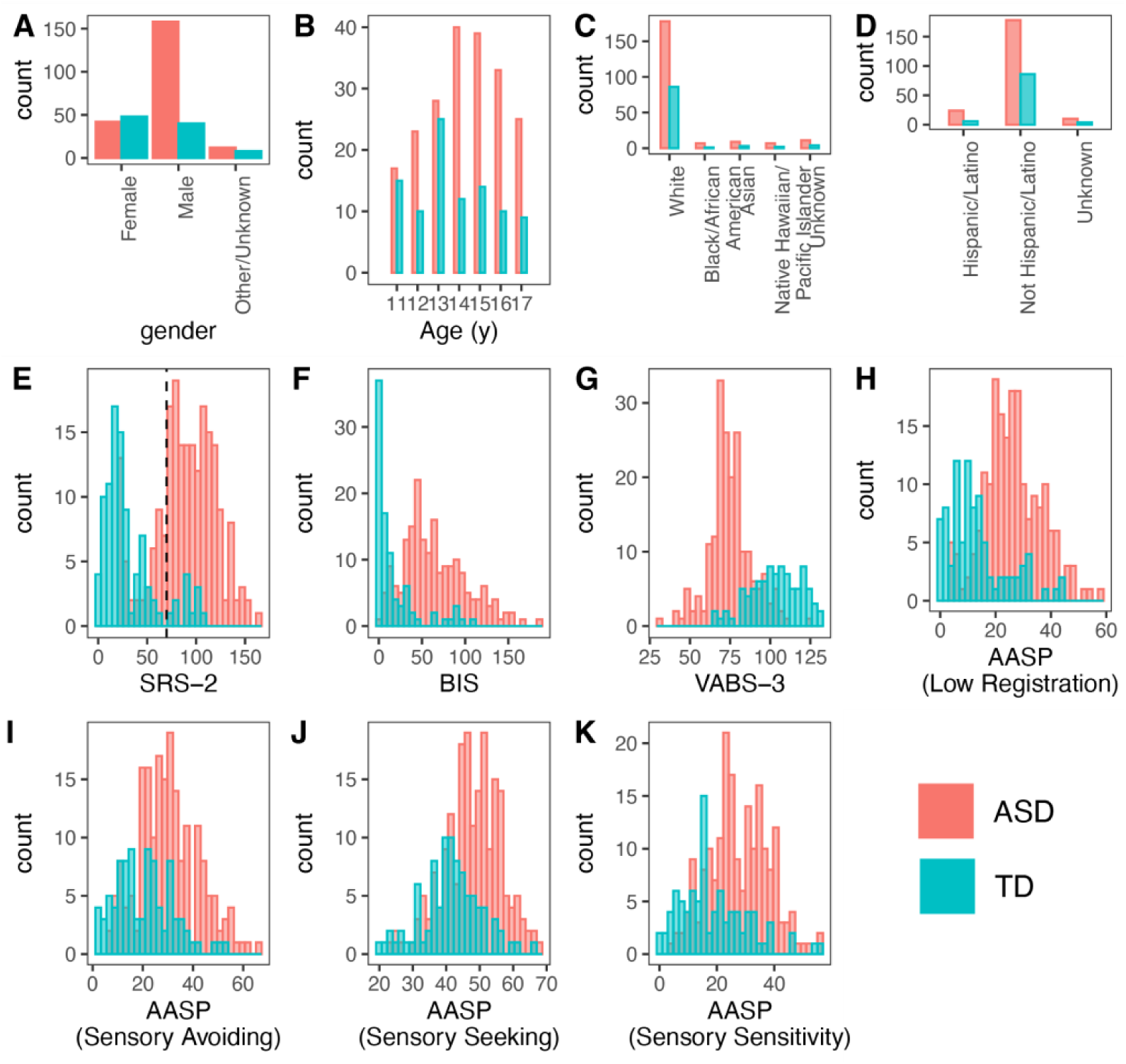
Participant demographics. Demographic distributions of our data indicate a large, heterogeneous sample. **A.** ASD participants included a higher proportion of males, as commonly noted in relevant literature. **B.** ASD and TD participants were evenly distributed in our selected age range (11 to 17 years). **C–D.** Distribution of race and ethnicity paralleled that of the US demographic. **E–G.** Distribution of SRS-2, BIS and VABS-3 scores suggested heterogeneity in symptom severity among ASD, also confirmed differences between TD and ASD. **F.** We also collected AASP, a questionnaire thought to assess detailed characteristics of sensory experiences in ASD. Note that VABS-3 and AASP (sensory seeking) scores were reverse scored to keep trends between ASD and TD consistent across all surveys.

**Supplementary Figure 3.**
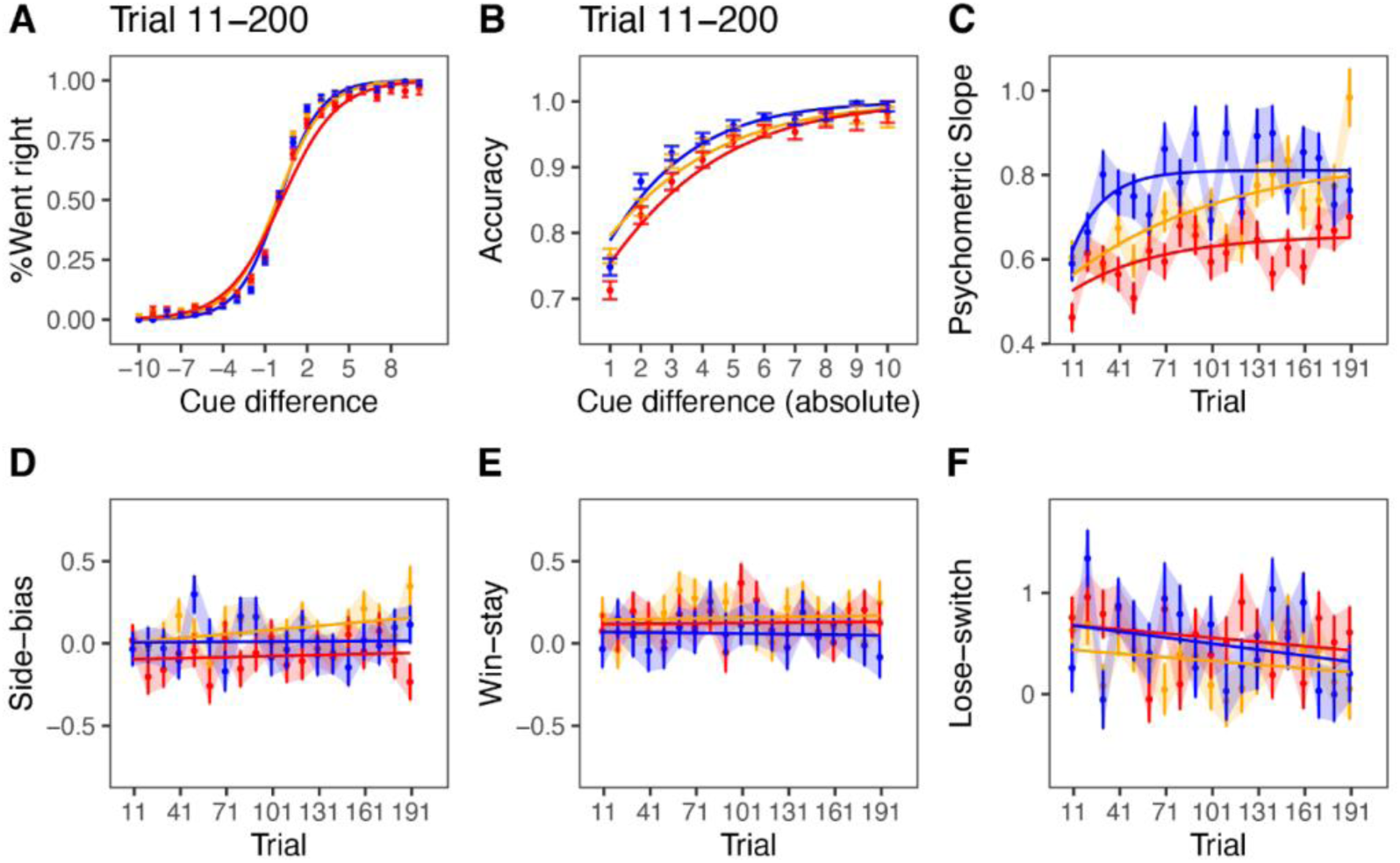
Psychometric performance. A–B. Psychometric function plots participants’ side choices (left or right) as a function of the flash or cue difference (signed and absolute), separately for TD (blue), ASD (gold), and ASD with lower VABS-3 scores (red). **C.** Variation in the slope of the psychometric with time in the task, estimated with a GLM fit to pooled data from each subject group and by taking bins of 10 consecutive trials. The GLM estimated choice probability as a function of flash difference (psychometric slope), win-stay and lose-switch biases, the intercept term captured side bias. Solid lines are learning curves fit to the model weights for flash difference. **D–F.** Variation in side-bias, win-stay, and lose-switch bias over time, positive and negative values indicate bias in favor of the right and left sides, respectively. Solid lines are linear regression models fit to the model weights. All error bars are SEM. All panels reflect participants who met task criteria.

**Supplementary Figure 4.**
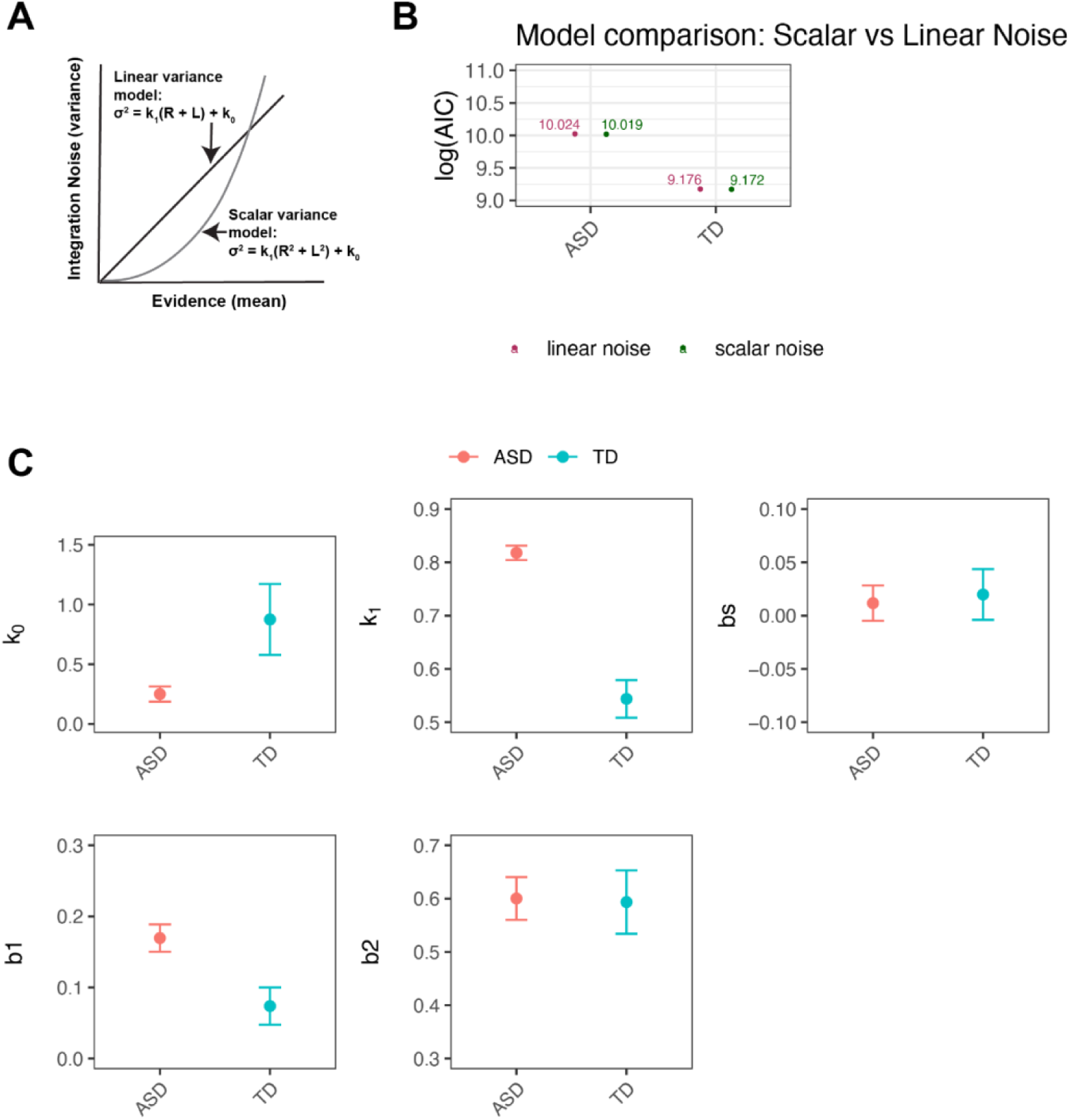
Linear vs Scalar variability signal detection theory models. **A.** Signal detection theory (SDT) models with linear and scalar noise. **B.** Comparison of the linear and scalar models for ASD and TD data. **C.** Model parameters for the linear noise model for ASD and TD.

**Supplementary Figure 5.**
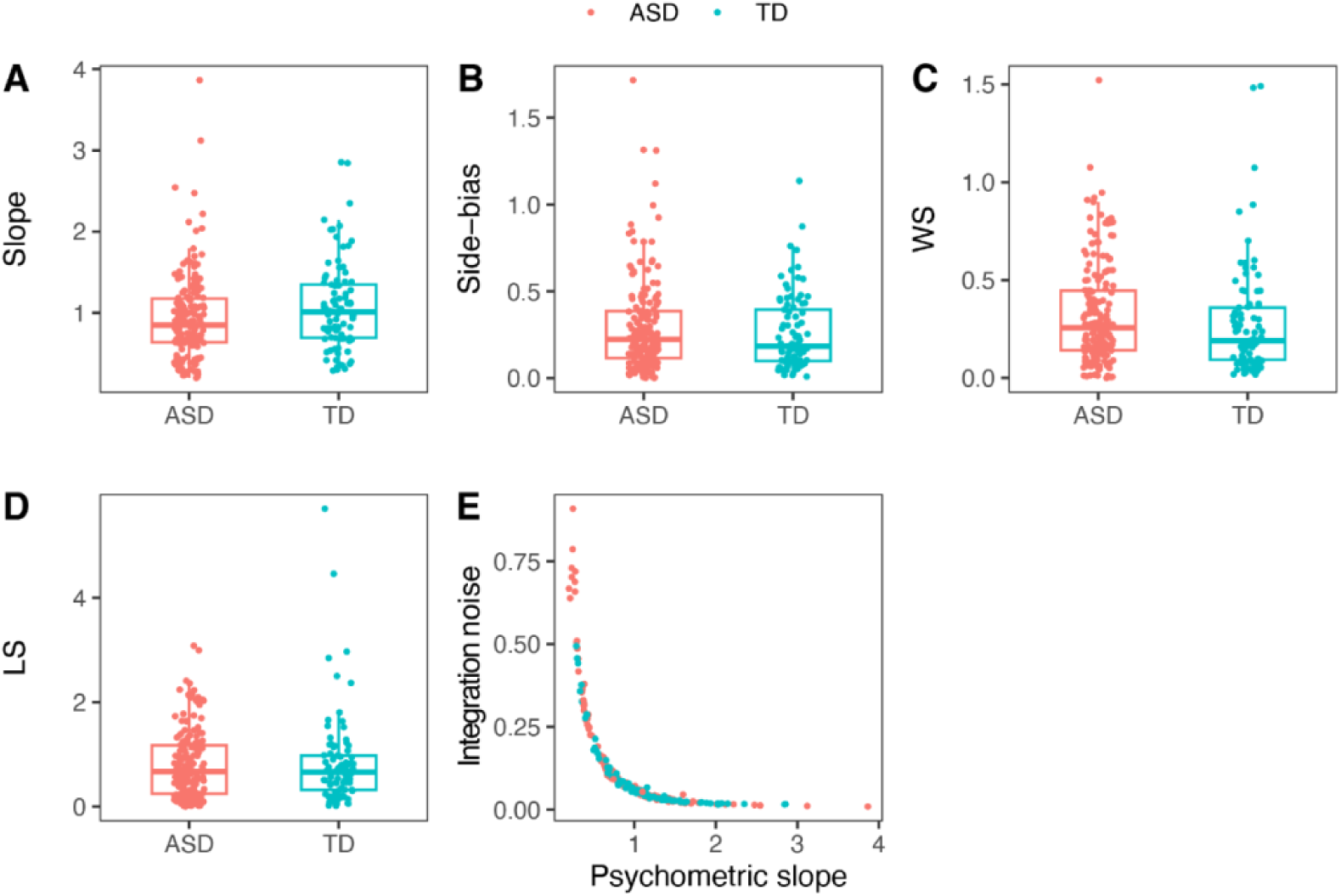
Individual participant psychometrics. A–D. We fit GLMs to individual participant data to estimate the psychometric slope, along with side bias, and trial history biases such as win-stay or lose-switch **E.** Correspondence between the psychometric slope estimated with GLM and integration noise (𝑘1) estimated with HBM-SDT.

**Supplemental Figure 6.**
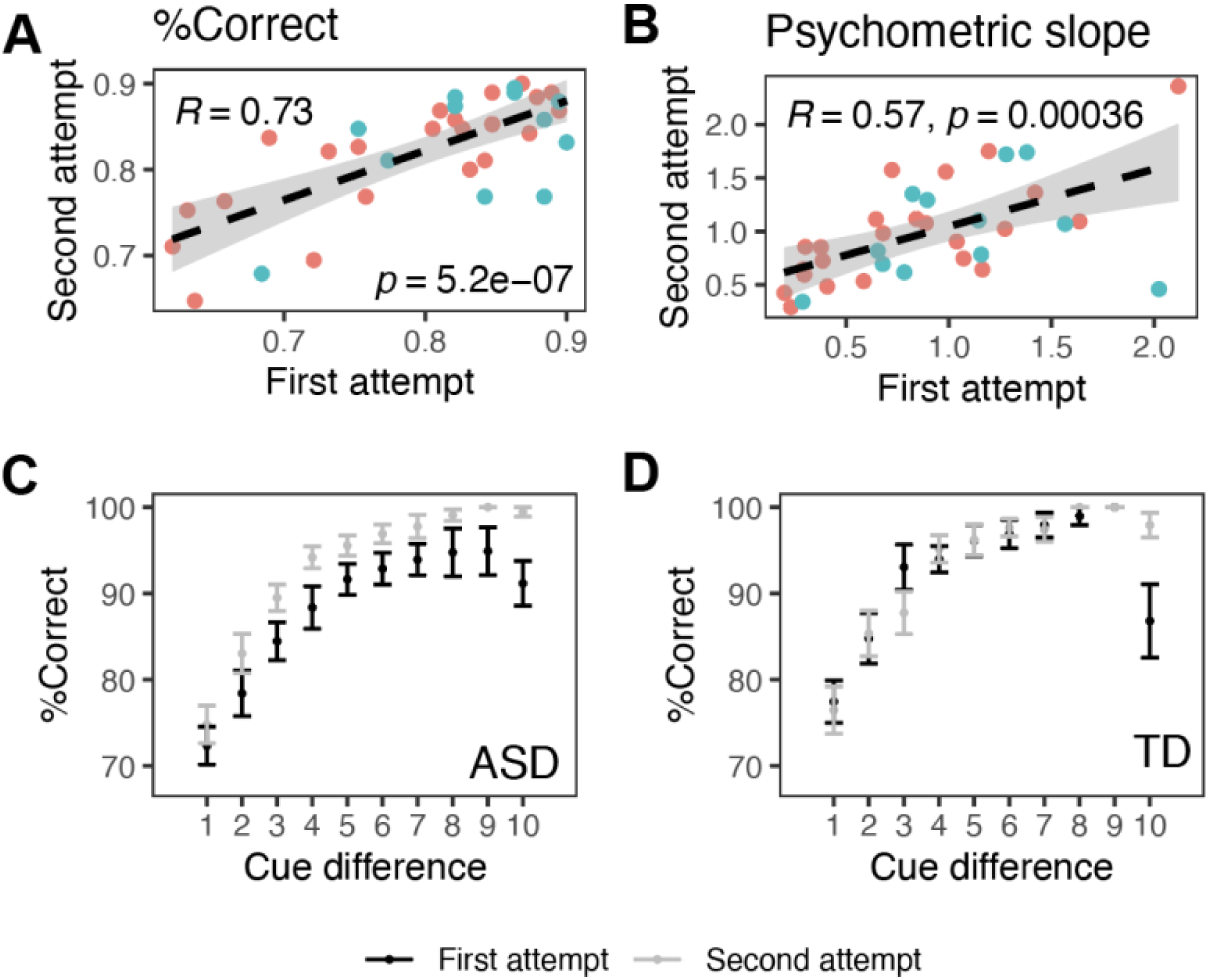
R**e**test **reliability.** 35 individuals (23 ASD; 12 TD) returned for a second session to estimate the retest reliability of the game. Accuracy (A) and the slope of the psychometric function (B) were highly correlated between the first and second attempt (P<0.001). Colored dots represent individual players, ASD in red, TD in blue. C) For the ASD cohort, accuracy in the second attempt was greater than the first. D) Psychometric performance across for the TD cohort was similar across first and second attempts.

## Notes

### Competing Interest Statement

The authors have declared no competing interest.

